# Non-linear Age-related Change in Human Interleukin-11 and the receptor subunit alpha DNA Methylation

**DOI:** 10.1101/2025.04.01.646733

**Authors:** Akiyoshi Shimura, Varun B Dwaraka, Kyosuke Yamanishi, Tomoteru Seki, Tsuyoshi Nishiguchi, Bun Aoyama, Takaya Ishii, Nathan James Phuong, Nipun Gorantla, Hieu Dinh Nguyen, Therese Santiago, Shota Nishitani, Ryan Smith, Gen Shinozaki

## Abstract

**Introduction:** Interleukin-11 (IL-11) is a cytokine involved in inflammatory processes and a previous study showed that blocking or knocking down *IL11* in mice prolongs a healthy lifespan. This study investigates DNA methylation (DNAm) changes in the *IL11* and IL-11 receptor subunit alpha (IL-11RA) gene across ages to reveal how aging might influence IL-11 production and sensitivity.

**Methods:** A genome-wide DNAm database focusing on Cytosine-phosphate-Guanine (CpG) sites within the *IL11* and *IL11RA* was analyzed. Hierarchical regression analyses examined the relationship between DNAm, age, and the squared age term for quadratic associations.

**Results:** The database comprised 10,297 samples (5,156 males and 5,141 females) with a mean age of 53.9 years (SD = 14.1 years). The majority of *IL11* and *IL11RA* CpG sites in the TSS1500 and 3’UTR regions exhibited significant inverse U-shaped associations with age. DNAm levels were low during youth, increased in middle age (40s-50s), and decreased again in older age.

**Conclusion:** The observed inverse U-shaped DNAm patterns in the *IL11* and *IL11RA* suggest n non-linear, age-related regulation of IL-11 expression and sensitivity. These findings indicate that IL-11 may have different roles across life stages and suggest that therapeutic interventions targeting IL-11 should consider age-specific effects.

## Introduction

Interleukin-11 (IL-11), also referred to as adipogenesis inhibitory factor (AGIF) and oprelvekin, is a pro-inflammatory and pro-fibrotic member of the IL-6 family [1]. IL-11 is associated with aging related processes. A recent animal model study [2] revealed that mice with knockout of the IL-11 gene (*IL11*) or IL-11 receptor subunit alpha (IL-11RA) gene (*IL11RA*) are protected against metabolic decline, multi-morbidity, and frailty in old age; moreover, administering IL-11 antibodies to aged mice improves metabolism and muscle function, reduces aging biomarkers, and extends lifespan by 20%. It has also been indicated that IL-11 signaling mediates senescence-associated pulmonary fibrosis in premature sesnescence model mice [3]. In humans, IL-11 levels are approximately 1.5 times higher in the centenarians than in middle-aged individuals [4]. Various fibro-inflammatory diseases such as rheumatoid arthritis, pulmonary fibrosis, nonalcoholic steatohepatitis, and kidney disease are associated with the upregulation of IL-11 [5]. A therapeutic approach targeting this pathway, such as the administration of IL-11 antibodies in acute kidney injury, not only alleviates inflammation and fibrosis but also restores kidney mass and function [6]. These findings highlight the potential of targeting IL-11 to control inflammation and extend a healthy lifespan.

On the other hand, IL-11 has physiological roles and beneficial aspects for health. IL-11 knockout mice exhibit reduced bone density and impaired bone formation from around three months of age, suggesting the importance of IL-11 in normal development during youth [7]. In females, IL-11 is crucial for decidualization in the pregnancy process [8]. Lower IL-11 production has been observed in the endometrium of patients with primary infertility, indicating its significant role in reproduction [9]. IL-11 formulations have also been reported to recover platelet counts during leukemia treatment [10]. In addition, IL-11 might have neuroprotective and anti-inflammatory effects, promoting neurogenesis and axon remyelination [11]. Thus, the role of IL-11 can have both beneficial and detrimental effects on the body.

Despite the limited human pathological and physiological studies on IL-11, it appears to be essential for normal development and reproduction in the first half of life, while potentially negatively affecting a healthy lifespan in the latter half. Given that plasma IL-11 levels are elevated in older adults compared to those in middle age, it is suggested that mechanisms exist to regulate IL-11 production and sensitivity along with the aging process. One such mechanism may involve age-related changes in the DNA methylation (DNAm) of *IL11* and associated genes. Therefore, the aim of this study is to investigate how the DNAm of *IL11* and its receptor gene, *IL11RA*, changes with age, using a genome-wide DNAm database from over 10,000 individuals.

## Methods

### Dataset

The analyzed samples are from a fully anonymized, de-identified human DNAm database collected from individuals who explicitly provided consent for academic use of their data through the personalized epigenome status measurement service provided by TruDiagnostic Inc. (Lexington, KY, US) between 2020 and 2023. The data collectors, holders, and analysts are independent of each other. The database contains DNA genome wide methylation information from 10,420 individuals, encompassing 864,628 Cytosine-phosphate-Guanine (CpG) sites. The analysis of this dataset was approved by the Institute of Regenerative and Cellular Medicine Institutional Review Board (IRB Approval number: #IRCM-2023-369).

### Blood collection and DNAm scanning

Blood samples were self-collected by participants using a finger-prick blood collection device, which involved obtaining a small blood sample from the fingertip (lancet and capillary method). This was stabilized with a lysis buffer. The samples were shipped to the TruDiagnositic lab, where the DNA was extracted, purified, and quantified. A total of 500 ng of DNA from each sample underwent bisulfite conversion using the EZ DNA Methylation kit (Zymo Research, USA). The bisulfite-converted DNA samples were then allocated to designated wells on the Infinium HumanMethylationEPIC BeadChip (Illumina Inc, USA), where they were amplified, hybridized, stained, and imaged using the Illumina iScan SQ instrument for further analysis.

### Dataset creation

The Minfi R package [12] was used for the pre-processing of DNAm data. All samples were pre-processed together to remove batch effects. During sample quality control, samples with aberrant methylation levels and background signal levels (mean P-value of ≥ 0.05) were removed. Probes with background signals above the same threshold were also discarded. The DNAm values were further normalized using the Genome-wide Median Normalization (GMQN) and Beta Mixture Quantile (BMIQ) methods. Missing values were imputed using the k-nearest neighbors (KNN) algorithm. Additionally, the composition ratio of white blood cells (WBC) was estimated using EpiDISH [13, 14], a method that showed a high correlation (R^2^ > 0.96) with immune cell subsets measured by RNA-seq and flow cytometry [15].

## Statistical analyses

First, the overall DNAm status of CpG sites related to *IL11* (12 sites) and *IL11RA* (16 sites) was examined. There are two different subunits for IL-11Rα, IL-11Rα1 and IL-11Rα2; however, these two subunits are currently not distinguished in terms of CpG sites. Second, multiple regression analyses were conducted with the DNAm beta values (β values) as the dependent variables, and age, sex, and estimated WBC composition as covariates. Since immune cell compositions may affect the DNAm status of *IL11* and *IL11RA* [16, 17], the composition of WBC was consistently adjusted in all subsequent analyses. M values were also generated by converting β values with residualizing the effects of WBC composition. The threshold for statistical significance was set at a P-value of < 0.05. Sex of the samples was determined based on DNAm information of the sex chromosomes [18]. Third, when age is simply input as a continuous variable in the multiple regression equation, there is an assumption that there is a linear relationship between age and DNAm. However, there is no guarantee that this relationship is linear. Therefore, to consider the possibility that the DNAm may increase (form a peak) or decrease (form a trough) at certain ages, a hierarchical multiple regression analysis using a quadratic model for age was performed to account for potential peaks or troughs in DNAm for the insignificant CpGs in the linear model (Supplemental figure 1).

## Results

Within the total of 10,420 samples, the age data was missing in 123 samples. The remaining 10,297 samples (98.8%) were used for the analysis. The dataset comprised of 5,156 males (50.1%) and 5,141 females (49.9%). The mean age of the sample was 53.9 years (SD = 14.1 years). Age composition is shown in Supplemental figure 2.

*IL11* and *IL11RA* methylation trend changing pattern with aging is shown in Figure 1 and overall DNAm status of *IL11* and *IL11RA* is shown in Figure 2. The result of the hierarchical regression analysis is also shown in supplemental table 1. In the age-linear model, for *IL11*, out of 12 CpG sites, 4 sites (cg01975672, cg16481281, cg15744492, and cg13508283) showed a significant increase in DNAm levels with age, while 1 site (cg13114229) showed a significant decrease. For *IL11RA*, out of 16 CpG sites, 10 sites (cg06226724, cg09219038, cg25028832, cg02306492, cg03534770, cg21504624, cg01178535, cg07564915, cg02656580, and cg01141771) showed significant decreases in DNAm levels with age.

**Figure 1.**
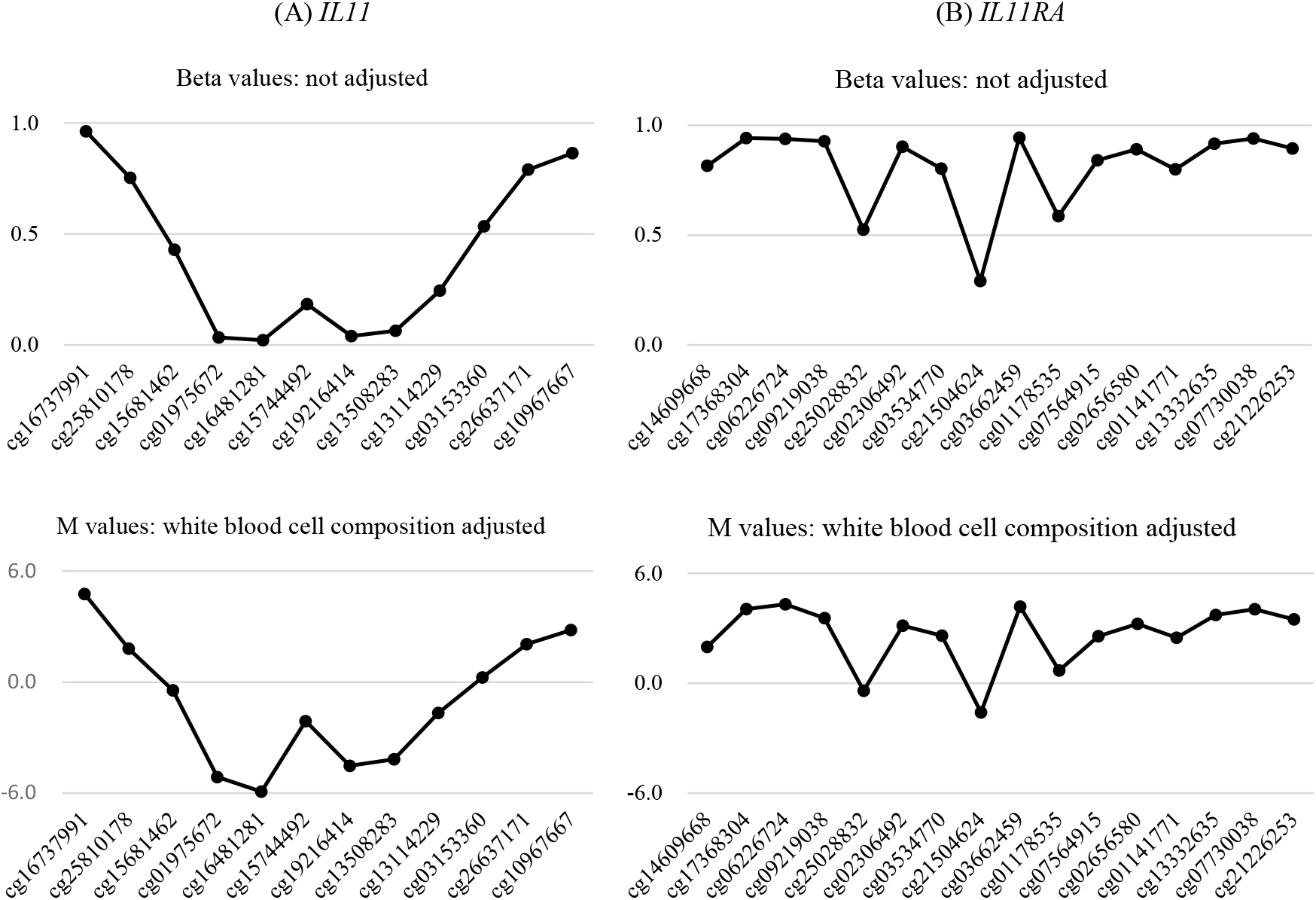
*IL-11 / IL11RA* methylation status. The upper graphs display the mean raw beta values for each CpG site, representing methylation status without adjustment for white blood cell (WBC) composition. The plots show the degree of methylation, where values close to 1 indicate complete methylation of the CpG site, and values close to 0 indicate complete demethylation. The lower graphs show the M values of the same CpG sites after regressing out the effects of WBC composition. Larger M values correspond to hyper methylation levels, whereas smaller M values indicate hypo methylation levels. Each CpG site is sorted according to its chromosomal position. In the IL-11 gene (A), the left side represents the 3’ UTR, while in the IL-11 RA gene (B), the right side represents the 3’ UTR.

**Figure 2.**
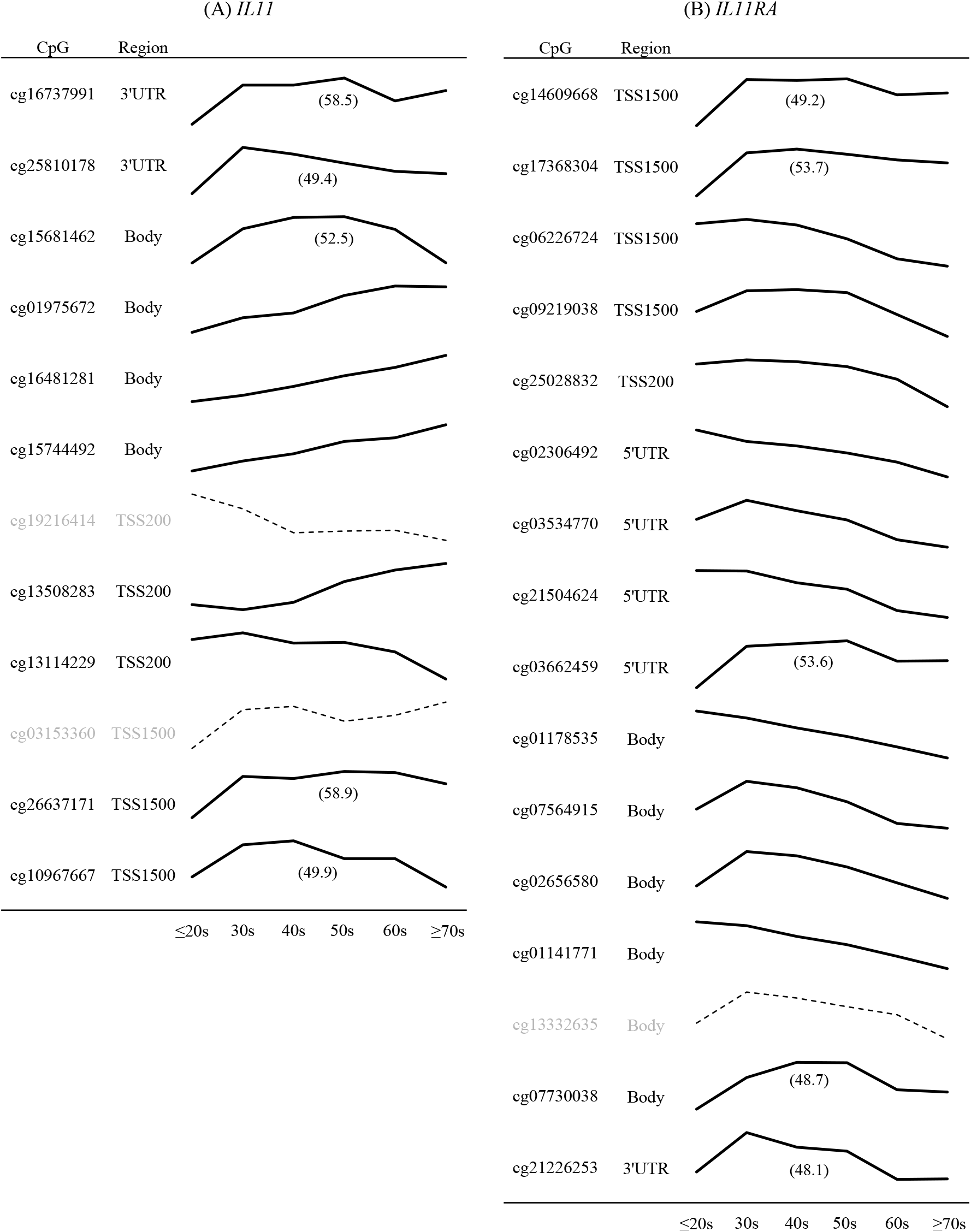
IL-11 / IL-11 RA gene methylation trend changing pattern with aging. Solid line graphs indicate significant linear or U-shaped associations with aging (P<0.05), while dashed line graphs indicate insignificant associations with aging. The values in parentheses represent the peak ages calculated by the quadratic regression model. The trend of each graph was adjusted by sex and white blood cell compositions, and the range of the graph is normalized.

In the age-quadratic model, for *IL11*, 5 out of 7 previously insignificant CpG sites (cg16737991, cg25810178, cg15681462, cg26637171, and cg10967667) showed significant inverse U-shaped associations with age, mainly in the TSS1500 and 3’UTR regions, with peaks between 49.4 and 58.9 years. The inclusion of the age^2^ term significantly improved R^2^, indicating non-linear DNAm regulation that increases until middle age and then declines. Similarly, for IL11RA, 5 out of 6 previously insignificant sites (cg14609668, cg17368304, cg03662459, cg07730038, and cg21226253) exhibited significant inverse U-shaped associations, with peaks between 48.1 and 53.7 years. The age^2^ term also significantly improved R^2^, showing DNAm decreases after middle age, especially in the TSS1500 and 3’UTR regions.

## Discussion

This study analyzed a large dataset collected from the general non-clinical population and found that the majority of the DNAm patterns in *IL11* and *IL11RA* do not change linearly with age. Instead, their DNAm levels increase towards middle age and then decrease as individuals become older. Notably, inverse U-shaped associations with aging were observed in the TSS1500 region, which is crucial for gene expression regulation, particularly repression [19], in both *IL11* and *IL11RA*, with peak ages between 48.1 and 58.9. This finding suggests that the IL-11 production potency as well as the sensitivity potency through the function of IL-11 receptor would be maintained at a young age, decrease towards middle age, and increase again beyond middle age. A previous study found increased IL-11 levels in centenarians compared to a middle-aged control group (mean age 49.0 years) [4]. This age of the control group coincides with the peak of the inverse U-shaped curve for its DNAm level found in this study, potentially amplifying the observed differences. An inverse U-shaped relationship with age, similar to that observed in the TSS1500 region, was also detected in the 3’UTR region. However, the role of the 3’UTR region is complex. It contributes to the regulation of messenger RNA localization, stability, and translation [20, 21], but the impact of its DNAm on gene expression, especially in *IL11* and *IL11RA*, is not well investigated.

IL-11 is involved in bone formation, remodeling, and maintaining normal trabecular bone mass [20]. IL-11 knockout mice exhibit significant bone density reduction from 3-4 months of age [7]. Therefore, IL-11 will need to be produced in the body during the active bone formation period at a young age. Additionally, as previously mentioned, IL-11 is essential for pregnancy process [8], suggesting that the IL-11 production potency must be maintained during the reproductive age. These physiological aspects are consistent with our findings that the DNAm level of *IL11* is lower at a young age. Conversely, the benefits of IL-11 in old age are not well-known. An increase in IL-11 production potency and sensitivity due to decreasing DNAm of *IL11* and *IL11RA* may cause heightened inflammation in the body and accelerate aging. In the future, approaches to intervene in the IL-11 pathway may be attempted for the prevention of various diseases associated with aging to extend a healthy lifespan. It is worth noting, however, that careful consideration is needed to target the appropriate starting age for interventions based on the data presented here.

This study has certain limitations. The non-clinical population sample used in this analysis was not collected through random sampling of the general population, which may introduce biases related to socio-economic factors or literacy. Additionally, racial and ethnic differences were not considered or adjusted for, and therefore, the generalizability of these findings to the broader population cannot be assured.

Future studies are needed to include large-scale measurements of IL-11 levels across different age groups in general populations to verify whether IL-11 levels and their sensitivity exhibit a trough during middle age and to assess the physiological significance of peak DNAm during this period. It would also be desirable to discover the mechanisms that cause the DNAm peak during middle age so that potential therapeutic interventions can be developed.

## Conclusion

The DNAm of *IL11* and *IL11RA* exhibits inverse U-shaped relationships with age, with their DNAm levels being low during youth, increasing during middle age (40s-50s), and decreasing again in older age, particularly around the TSS1500 region and the 3’UTR region. This pattern may reflect an age-related gene expression regulation process concerning the production of IL-11 and the sensitivity of its pathway. In the future, approaches to intervene in the IL-11 pathway might be attempted for extending a healthy lifespan; however, careful consideration is needed to target the appropriate age for interventions based on the findings that the gene regulation, especially by methylation, would differ between ages.

## Data availability

The data that support the findings of this study are not publicly available due to protection of patient data in accordance with maintaining HIPAA compliance. However, the data can be made available upon reasonable request after signing a Data Use Agreement.

## Acknowledgments

The dataset was provided free of charge by TruDiagnostic Inc. for this investigation. No external funding was received for this study.

---

**Supplemental figure 1.**
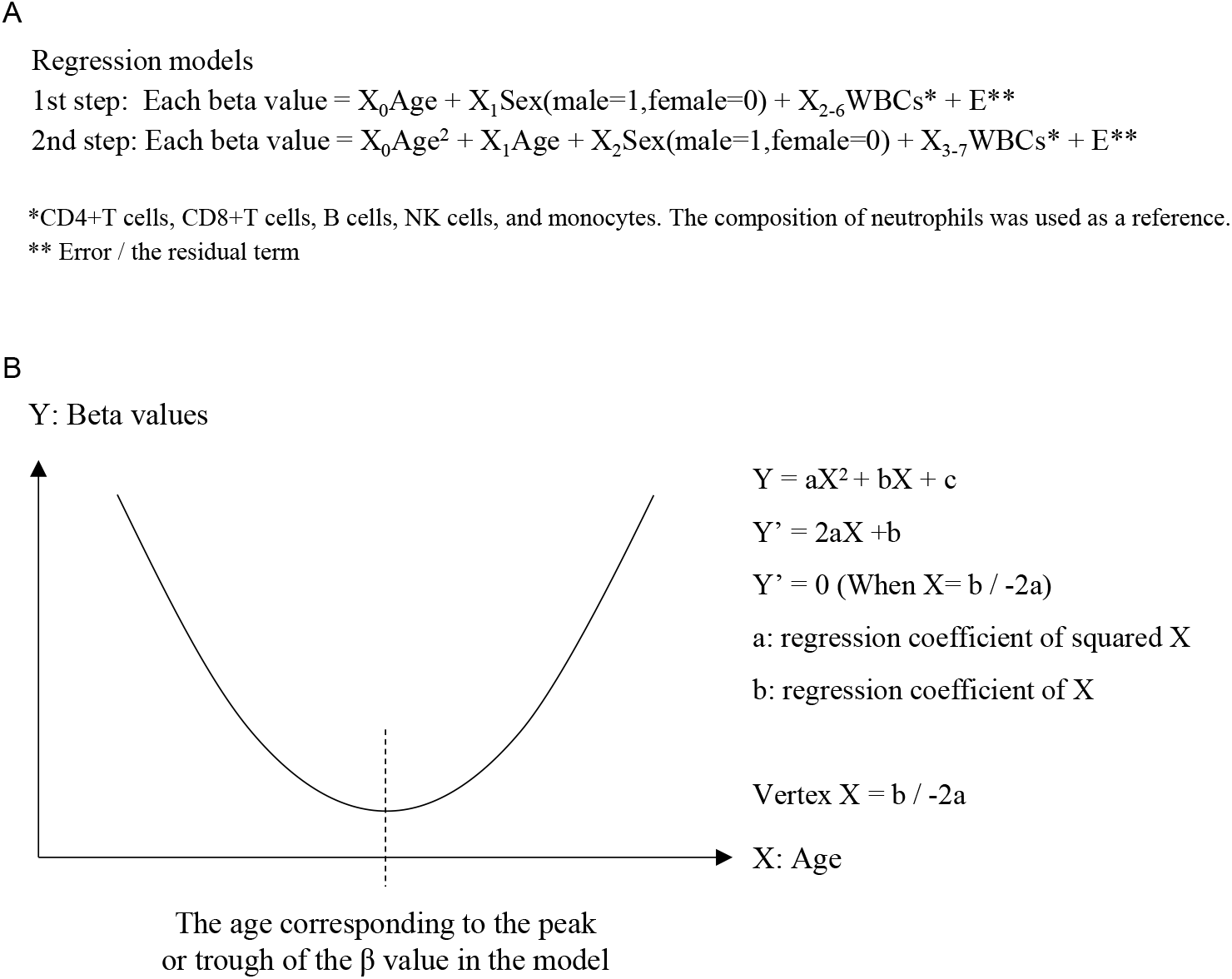
Explanation of quadratic equation model regression analysis. A: Explanation of the model equations used in the hierarchical multiple regression analysis. B: Calculation formula used to determine the peak age from the quadratic model.

**Supplemental figure 2.**
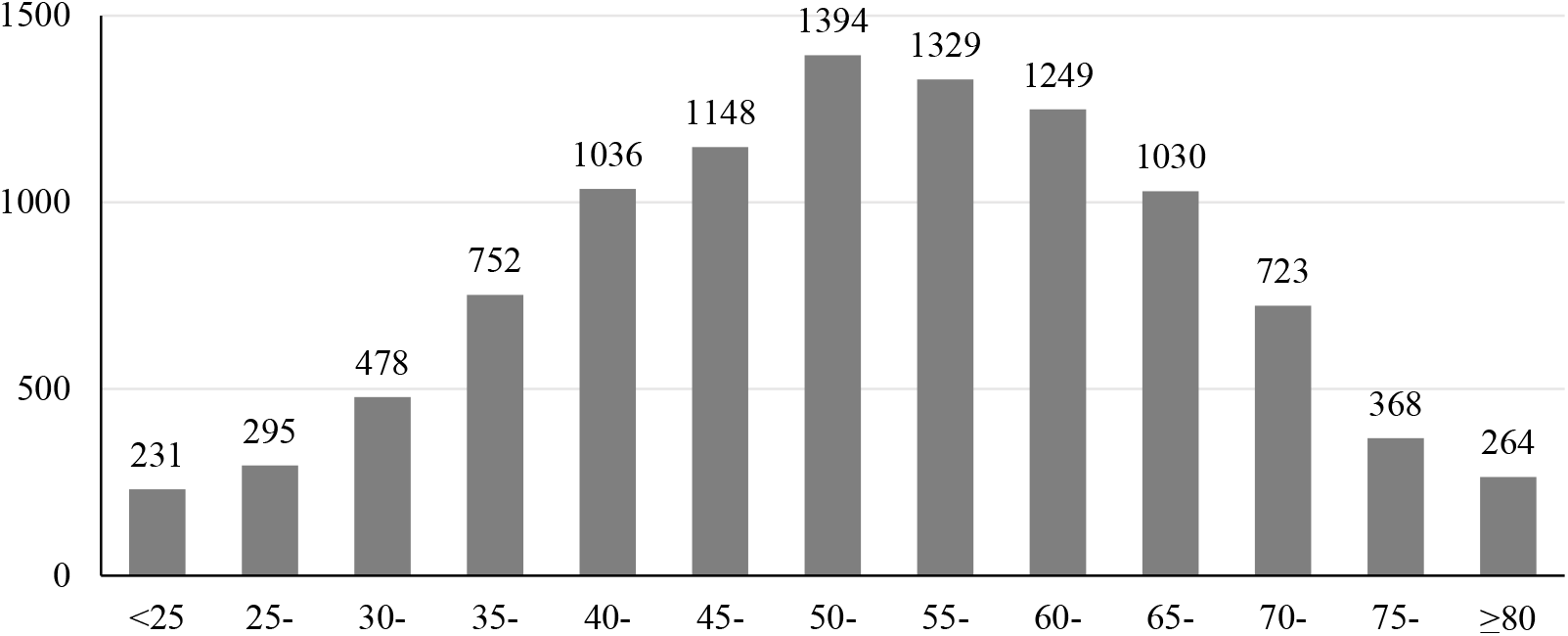
Age composition

**Supplemental table 1.**
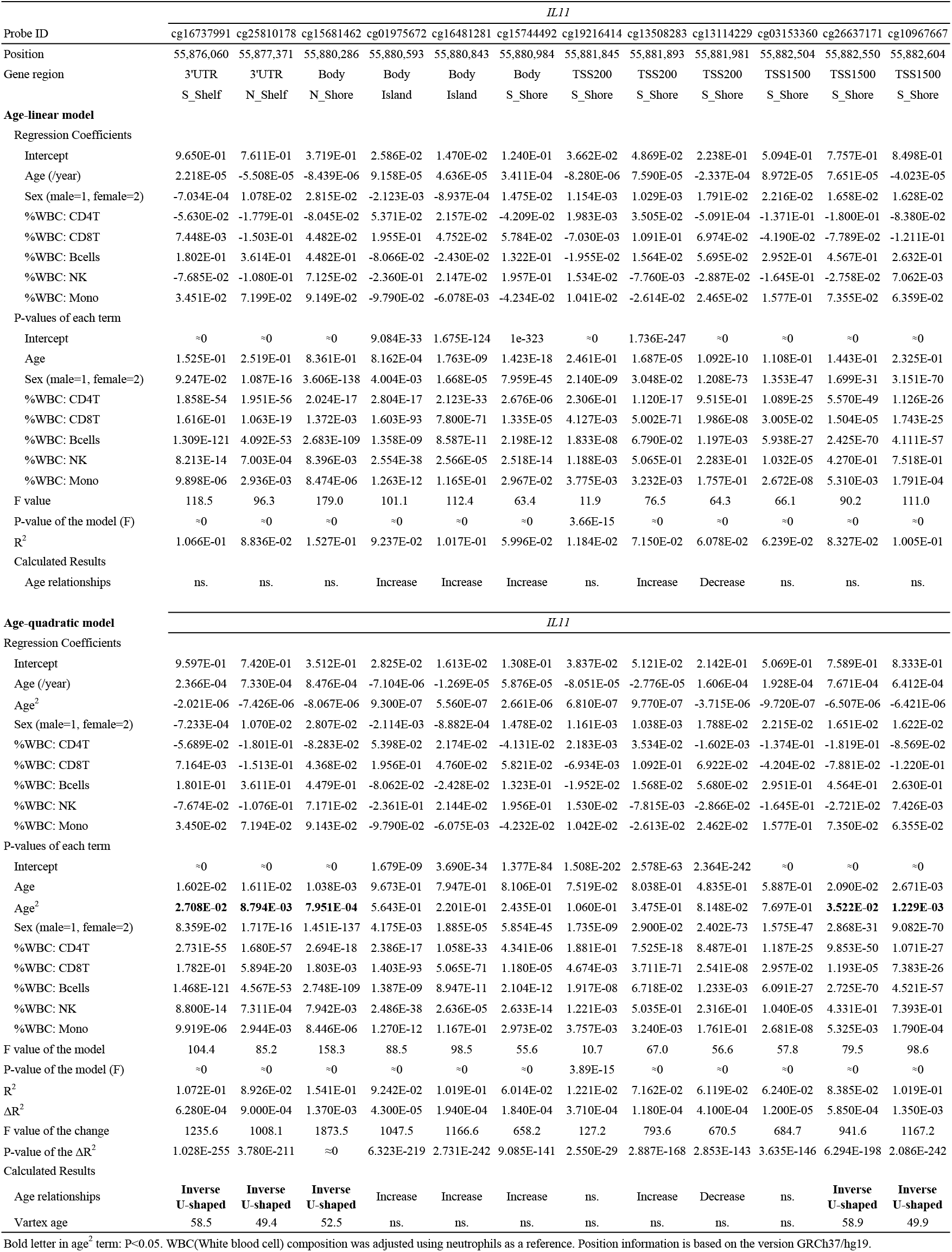
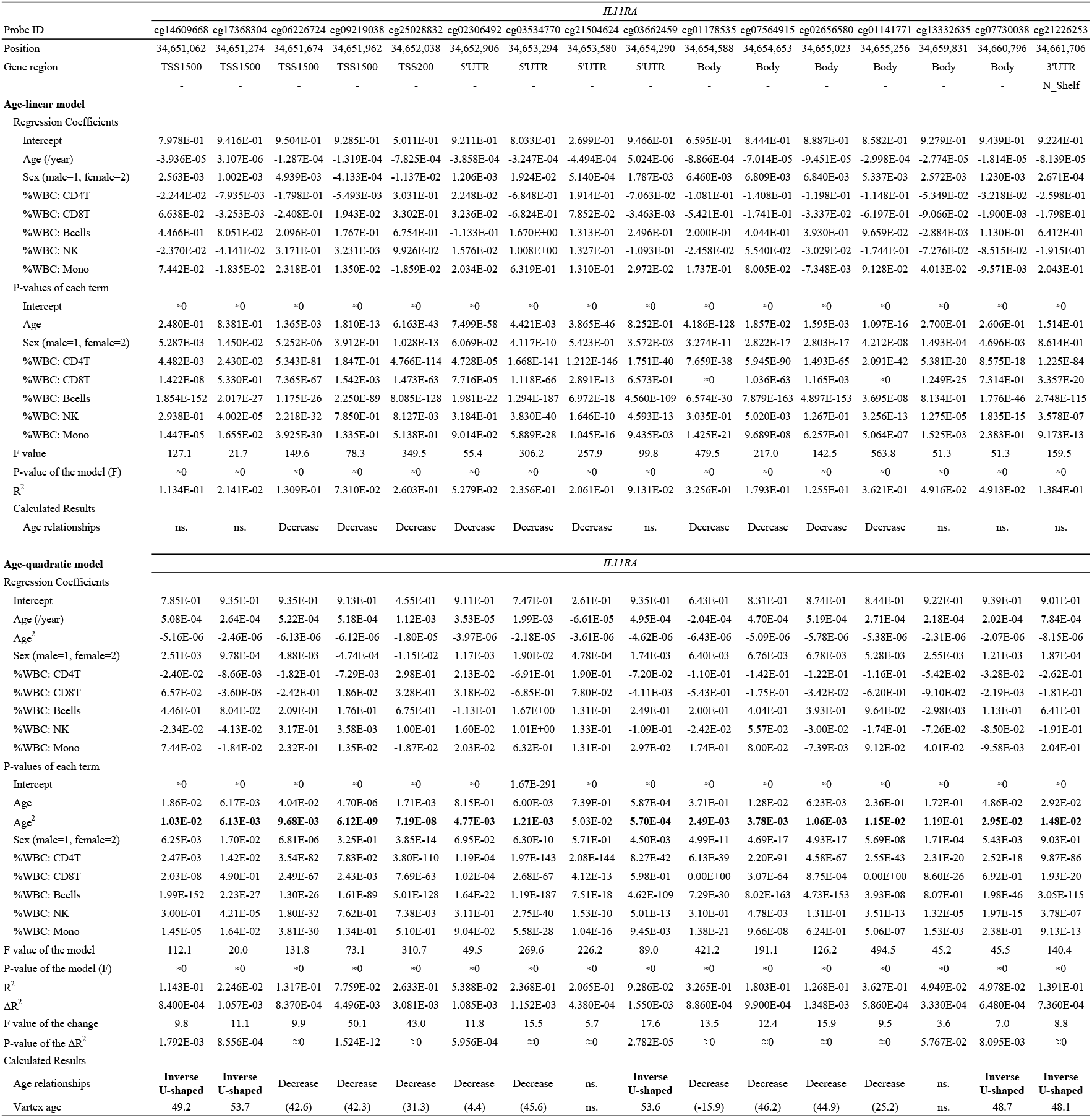
Human *IL11* and *IL11RA* methylated probes: the effect of age, sex, and write blood cells proportion on DNA methylation by multivariate regression analysis.

